# Olfaction foraging in visually oriented tropical arboreal ants *Oecophylla smaragdina*: Implications for insect predation studies using artificial sentinel prey

**DOI:** 10.1101/2023.11.14.566109

**Authors:** Lin Yan, Samuel Paul Kagame, Yang Liu, Takafumi Mizuno, Akihiro Nakamura

## Abstract

Predation is the key to understanding trophic interactions. Because of the brief and cryptic nature of predatory behavior, sentinel prey has been widely adopted as an indirect way to identify predators and understand trophic interactions. However, sentinel prey presents only static visual cues, potentially biasing toward visually oriented predators whilst ignoring those that require other sensory cues to forage. Despite this, the effectiveness of sentinel prey has rarely been tested. Here, we focused on the weaver ant, *Oecophylla smaragdina*, a keystone predator widespread in the Asian and Australian tropics. As this species has large eyes and is known to visually navigate in their arboreal habitats, we hypothesized that they rely on visual cues to forage and that their predatory behavior will be captured by caterpillar-shaped sentinel prey. Ants were collected as colonies, and preference trials on baits were conducted using combinations of olfactory and static visual cues including the caterpillar shape. Surprisingly, *O. smaragdina* showed little or no preference for baits in the absence of olfactory cues and did not differentiate the shapes of baits regardless of the presence of olfactory cues. Our results indicate that *O. smaragdina* is likely to make predatory decisions based primarily on olfactory cues, while visual cues might be used for other behaviors. Furthermore, *O. smaragdina* is likely to be left out by the predation studies using sentinel prey models which is particularly alarming considering the dominant role of this species in the trophic interactions of tropical rainforests. Our study demonstrates that morphological characteristics, arboreal habitats, and visually oriented behavior do not necessarily suggest the use of static visual cues for predatory decisions. We suggest that sentinel prey models should not be used alone when the dominant predators are unlikely to use visual cues to make predatory decisions.

## 1. Introduction

Predators are a key ecosystem component that regulates population dynamics and ecosystem functions through an evolutionary arms race between predators and their prey (Peckarsky & al. 2008). Other than directly consuming prey, predators can indirectly shape prey development, behavior, dispersion, and aggregations (Peckarsky & al. 2008, Lima 1998, Schmitz & al. 2004, Preisser & al. 2007). Predation on herbivores, one of the major sources of prey for many predators, indirectly affects plants and primary production through trophic cascading (Finke & Denno 2004, Finke & Denno 2005). Despite the ecological significance of predation, the cryptic and transient nature of predation poses a challenge to understanding its dynamics, especially at larger spatial and temporal scales (Howe & al. 2009, Crawley 2009).

Live prey can be used as bait to directly measure predation events; however, it is difficult to record the identity of the predator without intensive observation or video recordings (Friend 1995). Alternatively, the use of sentinel prey models, such as plasticine, molded to resemble prey, have been used to indirectly infer the coarse identity of the predator (birds, ants, wasps, and snails, etc.) using imprints left from attacks (Howe & al. 2009, Rößler & al. 2018). This has been widely adopted in ecological studies (Liu & al. 2020, Roslin & al. 2017), and the flexibility of this method has given rise to experimental manipulations of numerous visual cues related to finding prey (e.g. texture, color, shape) to understand the sensory ecology of the animals in question (Pan & al. 2021, Sam & al. 2015, Zvereva & al. 2019).

However, it is worth noting that although artificial prey may visually resemble prey (from a human perspective), it may underestimate predation compared with live prey that provide multiple cues including motion, chemical and tactile cues that may attract predators, and may overestimate predation because the lack of predator evasion mechanisms (Zvereva & al. 2019, Rößler & al. 2018, Zvereva & Kozlov 2023). For example, Roslin et al. (2017) concluded that arthropods are driving the high predation risk in the tropics using sentinel caterpillars. However, the finding could be biased by the possibility that more arthropod predators use static visual cues to make predatory decisions, and their method did not capture the predation events based on other cues.

Therefore, groups of predators that do not rely on static visual cues for prey recognition may be left out by the studies using this method, potentially leading to biased conclusions (Jackson & Pollard 1996, Anton & Gnatzy 1998, Nimalrathna & al. 2023). The sensory cues used by predators vary depending on the predator identities as well as the habitats where the interactions occur (Kielty & al. 1996, Ward & al. 2007, Short 2020). Thus the estimation of predation rate inferred from artificial sentinel prey may be highly variable among the habitats where predators may or may not utilize static visual cues to distinguish the shapes and colors of their prey. Of particular concern are invertebrate predators, which vary widely in the sensory cues used for predation compared with vertebrate predators that rely heavily on visual cues (Klärner & Barth 1982, Jackson & Pollard 1996, Wen & al. 2017). Furthermore, some invertebrates could be dominant predators, yet their key role in trophic interactions might be left out if predation is estimated using sentinel prey models. However, few studies have examined the foraging ecology of dominant species in a given ecosystem so that the potential bias introduced by the use of artificial sentinel prey could be corrected. Testing their response to artificial sentinel prey will give direct insights into the effectiveness of this method in understanding predation in tropical forest canopies.

Due to their ubiquitous occurrence and abundance, ants are regarded as one of the major predator groups, especially in the tropical regions (Jeanne 1979, Schmitz & al. 2000, Floren & al. 2002). As artificial sentinel prey are generally placed on leaves and stems of plants (Howe & al. 2009), marks left by ants can be attributed to those with arboreal affinities. While several studies concluded that ants were responsible for a large portion of attacks on plasticine caterpillars (e.g., Liu & al. 2020, Tvardikova & Novotny 2012, Leles & al. 2017, Tiede & al. 2017), their conclusions were based on the bite marks on the plasticine models and did not directly observe ants attacking them. A recent study by Zvereva & Kozlov (2023) found that arboreal ants did not attack plasticine caterpillars in boreal forests. This leaves a question as to whether arboreal ants of the tropical regions, especially dominant species such as weaver ants, *Oecophylla smaragdina*, attack plasticine models. *Oecophylla smaragdina* is one of the dominant species in the Indo-Australian tropics, and is known as one of the most active predators in the agricultural and natural systems of this region (Forbes & Northfield 2017, Tsuji & al. 2004). If plasticine models fail to capture predation by this and other dominant arboreal ant species, it has serious implications not only for ecological studies but also for the management of pest controls in production lands where plasticine models are used to monitor the effectiveness of predation by *O. smaragdina* (Denan & al. 2023).

In addition to examining the effectiveness of sentinel prey, we use this method to examine the sensory ecology of *O. smaragdina*, in the context of foraging. Several pieces of evidence have indirectly suggested that, unlike most ant species that rely on olfactory cues, *O. smaragdina* relies on visual cues for foraging (Mishra & Bhadani 2017, Lokkers 1990), although was not tested with behavioral trials but rather indirectly. Studies have demonstrated that their ability to discriminate patterns, well-developed eye morphology, and diurnal foraging lifestyle was correlated with the necessity of vision during foraging (Mishra & Bhadani 2017, Lokkers 1990). The arboreal nature of this species also corroborates with their visual capacity: canopy-dwelling organisms use visual cues due to better light conditions and rapid evaporation of chemical cues in the canopy (van Oudenhove & al. 2011, Loiselle & Farji-Brener 2002, Short 2020). However, no experimental studies have been conducted to directly examine the utilization of visual cues in the food preference of *O. smaragdina* (but see Jander and Jander 1998 for role of visual cues in navigation during foraging).

In the present study, we aim to examine the effectiveness of the use of sentinel caterpillars in estimating predation rates with a case study of *Oecophylla smaragdina*. We employed manipulative experiments in controlled lab conditions using both edible baits and plasticine sentinel models to test the relative importance of olfactory and static visual cues for their predatory decisions, connecting the sensory ecology of the weaver ant with the effectiveness of sentinel caterpillar in capturing predation by them. Based on the previous studies on visual acuity and diurnal foraging of this species, we hypothesized that *O. smaragdina* prioritizes visual cues rather than olfactory cues for predation. Furthermore, because sentinel caterpillars are visually effective in resembling prey, they will be preferred visually by *O. smaragdina*, that is, artificial caterpillar models effectively capture their predatory behavior.

## 2. Materials and Methods

### 2.1 Ant Collections

Nests of *Oecophylla smaragdina* were collected in Mengla County, Xishuangbanna Dai Autonomous Prefecture, located in the monsoonal tropical region of southern Yunnan Province, China in November 2020. To explore the connection between the difference in ecological factors in a different habitat, and potential adaptation by *O. smaragdina* in different populations, we included ants from two distinct habitats. We visited two locations representative of the dominant habitats of *O. smaragdina*: a tropical rainforest in Bubeng (21.61N 101.57E) and a rubber plantation located within Xishuangbanna Tropical Botanical Garden (XTBG; 21° 55’ N, 101° 15’ E) in Menglun. A total of three leaf nests were collected from the rainforest canopies using a canopy crane (80 m high, 60 m jib length) established in Bubeng that allowed us to access canopy trees at various heights. All three ant nests were collected from one of the dominant canopy tree species in this region, *Parashorea chinensis*, at heights over 50 m above ground using a pruning knife. In the rubber plantation, we collected a total of three ant nests on rubber trees (*Hevea brasiliensis*) at approximately 5 to 10 meters high by cutting the nests from tree branches using a pruning knife. All nests were transferred, kept separately in plastic boxes, and provided with sugar water.

Ant nests from the rainforest were larger (130-300 mm in diameter) than those collected from the rubber plantation (110-240 mm in diameter). We therefore confirmed species identity using DNA barcodes. In addition, as relative eye size may be associated with ants’ visual capacity, we measured eye and head areas to assess whether relative eye size varied between ant nests and collecting localities. Please see the Supplementary Information for the details of DNA barcoding and eye size measures.

### 2.2 Experimental Design

All behavioral trials were conducted within 2 weeks of collecting of the colonies from the field. We designed four manipulative experiments to test: (1) the importance of visual shape in the absence of olfactory cues (the **Visual-only experiment: Fig. 1a**); (2) the importance of olfactory cues and natural prey (the **Olfaction and natural shape experiment: Fig.1b**); and (3) the interaction between visual and olfactory cues (the **Olfaction and caterpillar shape experiment**: **Fig.1c**); and (4) the effects of olfactory cues without the presence of visual cues (the **Darkroom experiment: Fig.1d**). All experiments were conducted within a plastic box with each ant colony placed in the middle, and four bait treatments at each of the four corners of the box (**Fig. 1**). Bait treatments were placed in transparent polypropylene plastic tubes.

**Figure 1.**
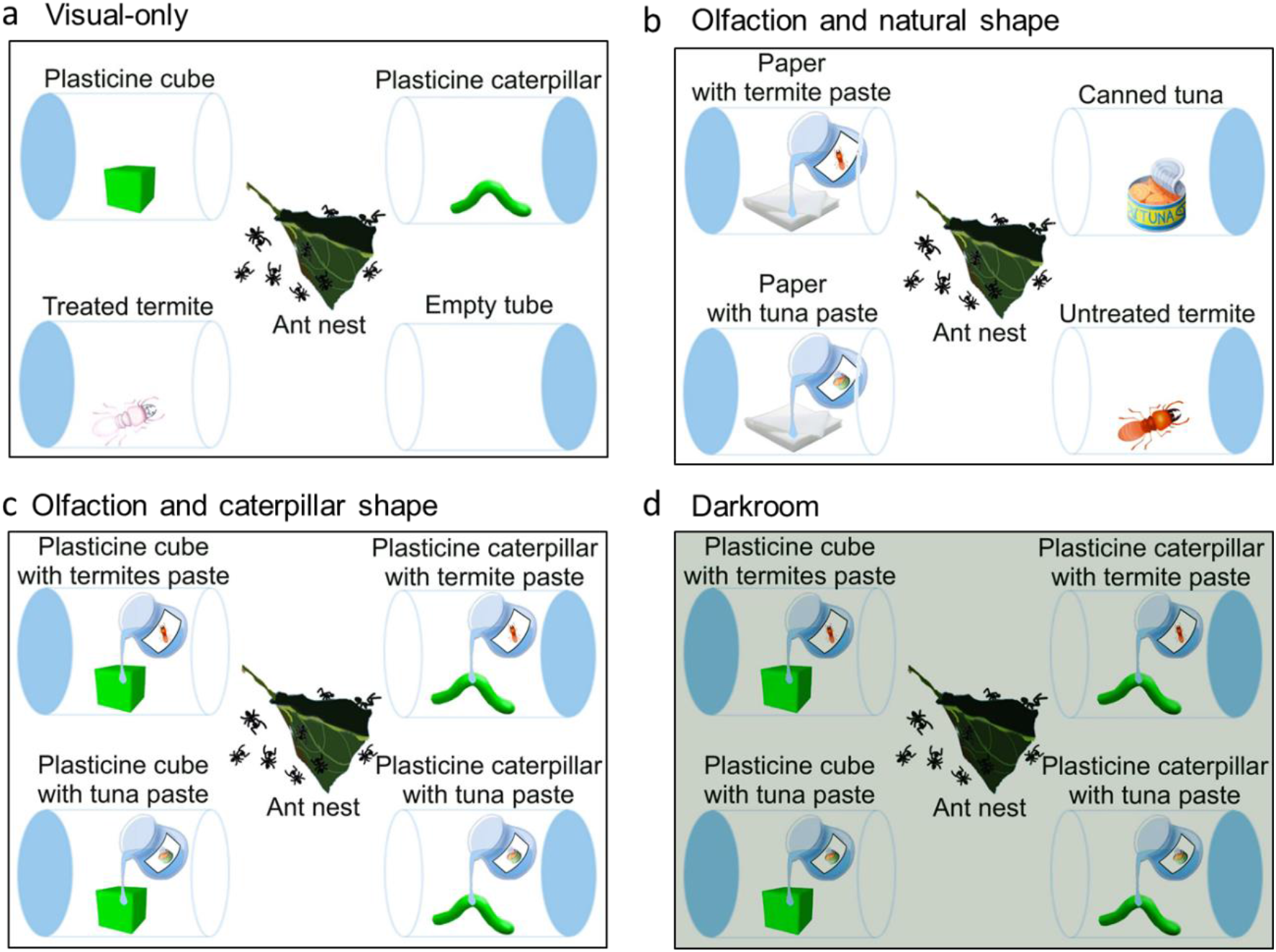
Conceptual illustrations depicting the design of the four experiments: **a.** the Visual-only experiment to assess the effect of visual cues; **b.** the Olfaction and natural shape experiment to test the effect of olfactory cues with the presence or absence of natural visual cues on the baits; **c.** the Olfaction and caterpillar shape experiment to test the interaction between olfactory cues and abstract visual cues of plasticine models; and **d.** the Darkroom experiment to test the effect of olfactory cues with the absence of visual cues.

As we had three small and three large nests collected from the two habitats respectively, we used a large box (550 mm ×370 mm ×380 mm depth) for larger nests and a small box (180 mm ×280 mm ×180 mm depth) for small nests. For bait treatments, we used large tubes (120 ml with 44 mm diameter and 105 mm depth) in the large boxes and small tubes (5 ml with 15 mm diameter and 49 mm depth) in the small boxes to keep the baits in them. Non-drying sticky glue (tanglefoot) was applied to the rim of the box (approximately 2 - 4 cm width) to prevent ants from escaping. Before the experiment, ants were starved (without sugar water) for at least 24 hours before the experiments began but provided with water. To quantify the ants’ preference for different types of bait, we recorded the number of ants presents inside each tube every 10 minutes for one hour. In addition, we also recorded the time when an ant first entered the tube.

#### 2.2.1 Visual cues

The Visual-only experiment (**Fig. 1a**) tested the effect of prey shape in the absence of olfactory cues using bait treatments consisting of (1) plasticine caterpillars, (2) plasticine cubes, (3) dead termites treated by critical point drying for minimized olfactory cues, and (4) an empty tube as a control. We used plasticine caterpillars, a classic model method widely used in ecological and behavioral studies (Rößler, Pröhl, and Lötters 2018), and plasticine cubes which differ from plasticine caterpillars by shape only. To construct plasticine baits, we used non-toxic green plasticine (Newplast, Newclay Products Limited, Newton Abbot, UK). Plasticine caterpillars (3 mm diameter ×30 mm length) were molded using a metal syringe (a syringe used for pastry molding), and the same quantity of plasticine was molded into cubes (7mm length) by hand. We selected the termite *Odontotermes yunnanensis* for use in our experiments. This termite species is a preferred prey species for *Oecophylla* under natural conditions in the study area (pers. obs.). Termites were collected from nests and only worker termites were used. To minimize olfactory cues from termites, we prepared specimens by critical point drying and processed by Leica EM CPD300 (Leica Microsystems GmbH, Wetzlar, Germany). The program was set as follows: cooling at 15 degrees Celsius, slow CO_2_ admittance with 120 seconds of delay, 16 exchange cycles with the speed set to “6”, followed by a slow heating process at 40 degrees Celsius, and finally a slow gas discharge. Supercritical drying has been used for chemical extraction as well as effective chemical removal in several fields (Maheshwari & al. 1995, Brunner 2010). For large nests, we placed three sentinel models (caterpillars or cubes) or 30 termites in each tube. For small nests, we used one sentinel model (caterpillars or cubes) and 10 treated termites.

#### 2.2.2 Olfactory and visual (natural shape) cues

The Olfaction and natural shape experiment (**Fig. 1b**) tested the importance of visual and olfactory cues combined using bait treatments consisting of (1) termite olfactory cues only, (2) tuna olfactory cues only, (3) termite olfactory and visual cues, and (4) tuna olfactory and visual cues. We used canned tuna because is a preferred bait that is widely used in *Oecophylla* studies (Andersen 091992, Narendra & al. 2012), To prepare olfactory cue treatments, we prepared paste by crushing worker termites or canned tuna using a pestle and mortar. A small quantity of water was added while crushing the bait until it became fine and smooth. Next, this termite or tuna paste was applied to filter paper (15 mm ×15 mm) until saturated. Larger tubes contained three pieces of paper and smaller tubes contained one piece of paper. Termite olfactory and visual cues bait treatment consisted of freshly killed, untreated termites (30 in large tubes, 10 in small tubes). Tuna olfactory and visual cue treatment consisted of canned tuna (1.2g of tuna in large tubes, 0.4 g of tuna in small tubes).

##### 2.2.3 Olfactory and visual (caterpillar shape) cues

The Olfaction and caterpillar shape experiment (**Fig. 1c**) tested the importance of olfactory and visual cues of plasticine models using bait treatments consisting of (1) tuna paste applied to plasticine caterpillars. (2) tuna paste applied to plasticine cubes, (3) termite paste applied to plasticine caterpillars, and (4) termite paste applied to plasticine cubes. We applied the tuna or termite paste to the plasticine cubes and caterpillars until all surfaces were covered. The larger test tubes contained three sentinel models in each tube, while the smaller test tubes contained one sentinel model.

#### 2.4.4. Absence of visual cues

The Darkroom experiment (**Fig. 1d**) tested the importance of olfactory cues without the presence of visual cues. To be consistent, the experimental setups were the same as the Olfaction and caterpillar shape experiment (**Fig. 1c**) but conducted in a dark room to eliminate visual cues. We used an infrared camera (Sony FDR-AX60) placed approximately 50 cm above the experimental setups to observe ant behavior. As an important caveat to this experiment, the infrared cameras made it difficult to verify whether ants were inside (or on) the tube. For this experiment, we, therefore, counted the number of ants which appeared to be on the tubes in the infrared video clips. Given our imaging difficulties, we did not measure the time that the first ant entered the tube.

### 2.3. Statistical Analysis

Statistical analyses were performed using R software (version 4.1.2). We first used a linear mixed model with the identity of ant nests as a random factor, to test for differences in relative eye size between ant nests collected from the rainforest and rubber plantation. Forest type, major/minor worker, and the proximity from the nest (near/far) were included as candidate predictors. After confirming the normality of residuals, a Type II Wald chi-square test was performed to test for the significance of the predictors using the car package (Fox & Weisberg 2018). The best model was selected based on the lowest Akaike information criterion (AIC).

We next tested for the differences in preference (i.e., the number of ants in the experimental tubes) to different baits in each of our four experiments. A generalized linear mixed model was used, initially with Poisson distribution but overdispersion was found when fitting our data using the glmer() function in the package lme4 (Bates & al. 2007). As a result, we fitted our data with a generalized linear mix model using Template Model Builder (glmmTMB package, (Brooks, Kristensen, Van Benthem, & al. 2017)) with a negative binomial distribution with linear parameterization, to account for count data in the present study and to control for zero inflation (Hardin & al. 2007, Brooks, Kristensen, van Benthem, & al. 2017). The candidate models in each experiment included the four experimental treatments (tubes), time intervals, habitats (rainforest and rubber plantation from which ant nests were collected), and the interaction between time and treatment as fixed effects. All models were constructed with the identity of ant nests as a random factor, a single zero-inflation parameter applied to all observations, and the size of the nests as an offset in all candidate models (Bolker 2016). Nest size was estimated in volume (cm^3^) and this offset was included to relax the assumption that more individuals from larger nests forage and attack baits (Bolker 2016). The best model was selected based on the lowest AIC as well as a significant difference from the null model (Arnold 2010). After the best model was selected for each treatment group, a Type II Wald chi-square tests were performed to determine the significance of fixed factors using the car package (Fox & Weisberg 2018). Tukey’s Honest Significant Difference (HSD) test was conducted as a posthoc test to identify the pairwise significance between estimated marginal means of treatments (emmeans package, (Searle & al. 1980)). Model predictions were generated using the best model and the original dataset, with the offset colony size set to 8000 cm^3^. The predictions were used to plot the regression lines with the observed data in scatter points.

In addition to the number of ants in the experimental tubes, we also assessed the time when the first ant entered the experimental tubes. Because the nature of the data violated the assumptions of a linear model, we adopted a non-parametric test (Kruskal-Wallis Rank Sum Tests) to compare the time when ants first arrived among the four types of baits in each experiment.

## 3. Results

### 3.1. Phylogenetic Analysis and Relative Eye Size

The ants from the two different habitats formed a monophyletic clade (**Fig. S1**) together with other specimens collected within the same locality and Thailand. *Oecophylla smaragdina* collected from Australia and India formed a sister group to our specimens.

The best model that explained relative eye size contained only major/minor worker as the predictor (χ2=17.63, df=1, p<0.001) and forest type was not in the best model indicating that changes in habitat types (rainforest and rubber plantation) did not influence the relative eye size of *O. smaragdina* **(Fig. S2a)**. Minor workers had a larger relative eye size compared to major workers (**Fig. S2b**).

### 3.2 Visual cues

The estimated marginal mean of the number of ants in the empty tubes was the highest (estimated marginal mean±SE = 2.12±0.98), followed by the tubes with treated termites (1.85±0.84), plasticine cubes (1.68±0.77), and plasticine caterpillars (1.23±0.57). The best model included the experimental treatment which was marginally significant (χ^2^=8.054, df=3, p=0.045). Posthoc analysis suggested differences between the empty tube and the tube with plasticine caterpillar (p =0.038). However, no significant difference was found in other pairs, including the plasticine caterpillar and cube (p=0.344), and between treated termites and empty tube (p=0.884) **(Table S1)**. The effect of time was excluded from the best model, as the number of ants visiting the tubes did not change over time **(Fig. 2a).** No significant difference was found in the time when an ant first entered the experimental tubes (Kruskal-Wallis Rank Sum Tests, χ^2^=1.384, df=3, p=0.709) **(Fig. 3a)**.

**Figure 2.**
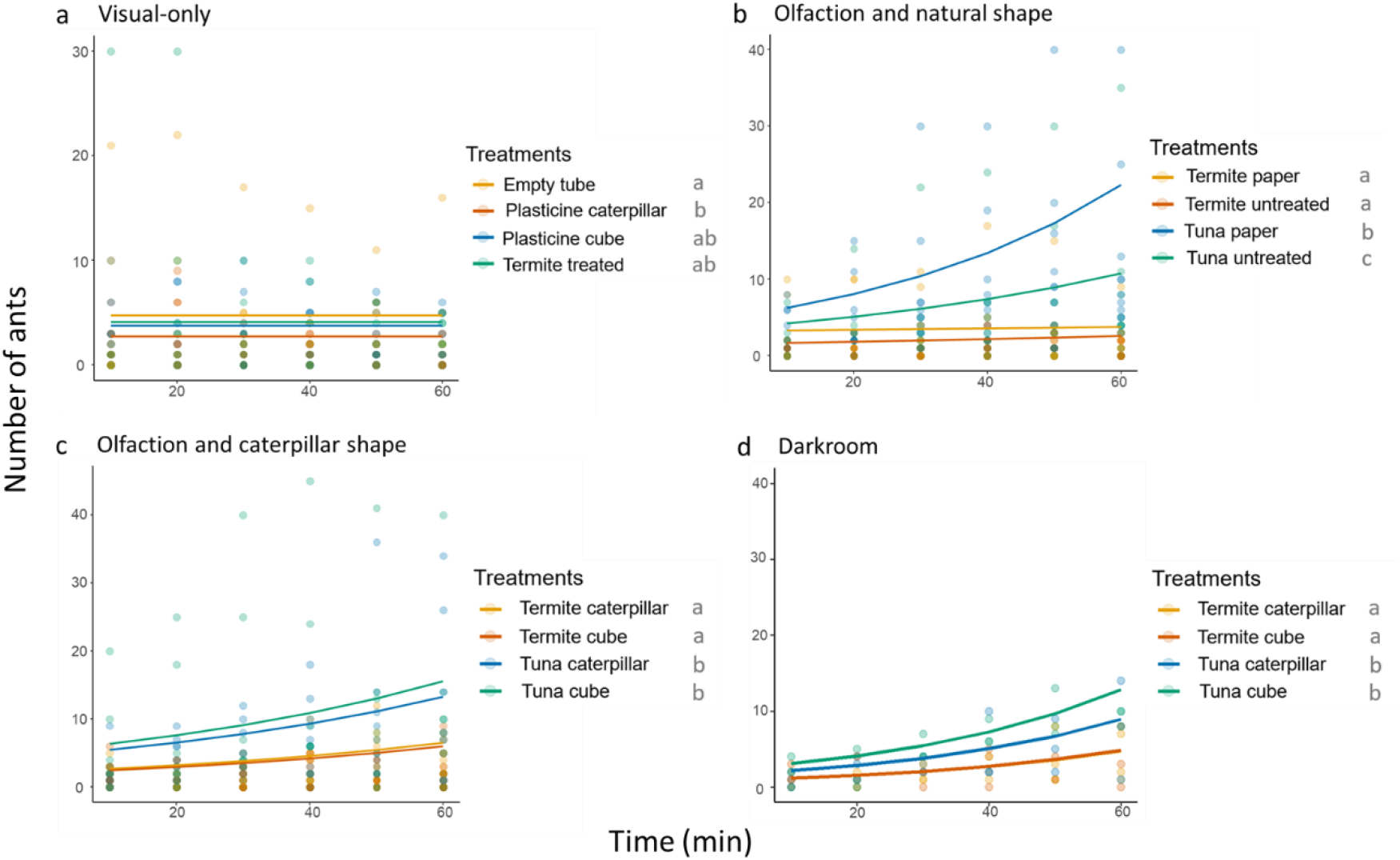
The number of ants found in each of the four experimental tubes over time (10- minute intervals) in: **a.** the Visual-only experiment; **b.** the Olfaction and natural shape experiment; **c.** the Olfaction and caterpillar shape experiment; and **d.** the Darkroom experiment. All regression lines are predictions based on the best model. Posthoc pairwise comparisons of the four experimental treatments are denoted by the letters beside each legend of the experiments (p<0.05, based on Tukey’s HSD test of estimated marginal means).

**Figure 3.**
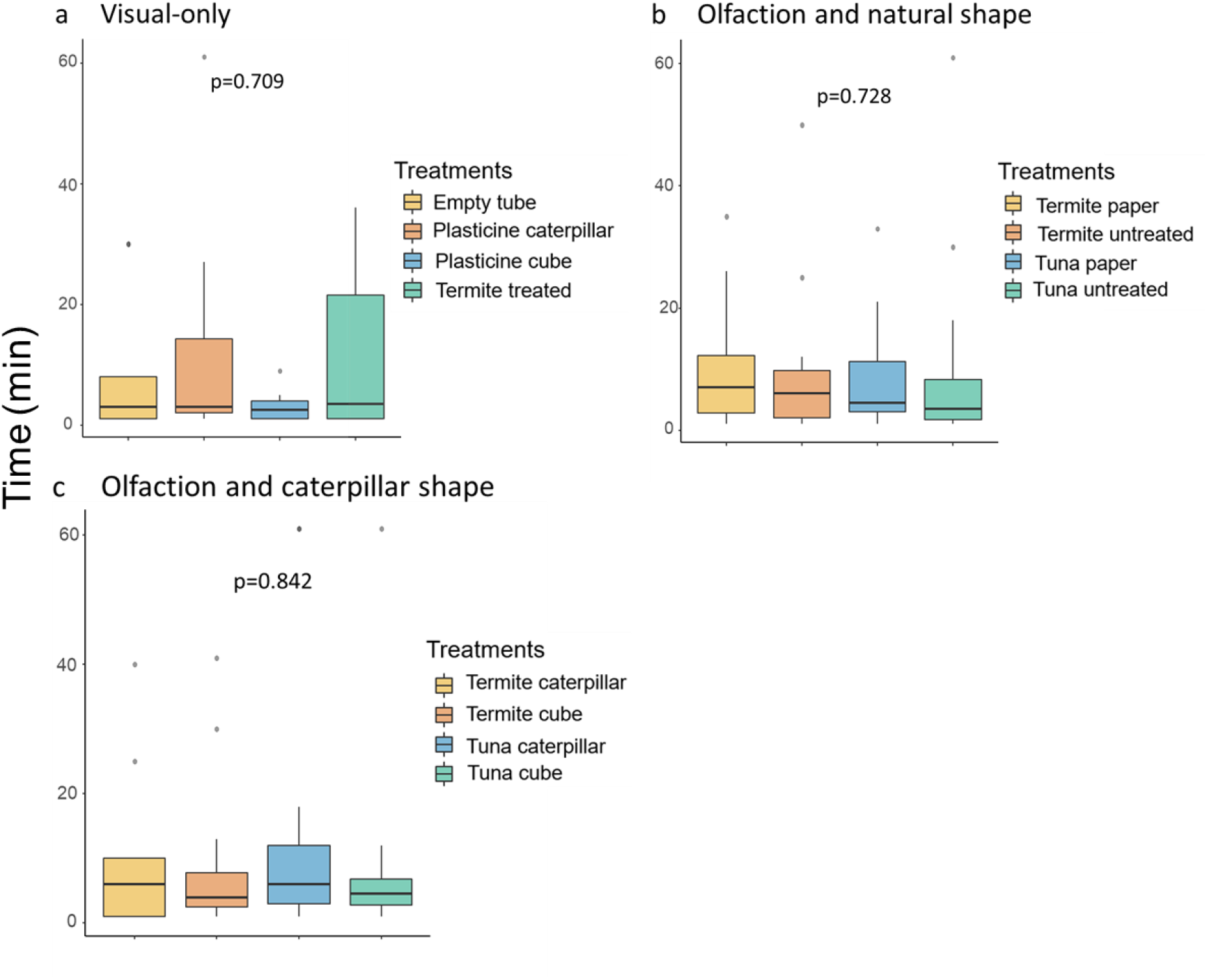
Box plots showing the time of first entry in minutes by an ant to the experimental tube in: **a.** the Visual-only experiment; **b.** the Olfaction and natural shape experiment; and **c.** the Olfaction and caterpillar shape experiment; The p-values from Kruskal-Wallis Rank Sum Tests are shown. As the infrared camera was not clear enough to confirm whether ants were inside or outside of the tubes, we did not test this for the Darkroom experiment.

### 3.3 Olfactory and visual (natural shape) cues

The largest number of ants was observed on papers with tuna paste (5.72±1.93), followed by tuna (3.09±1.08). Termite paper was visited less frequently (1.50±0.55) and untreated termites was hardly visited by ants (0.94±0.36) **(Fig. 2b)**. The best model included the bait type (χ^2^=123.03, df=3, p<0.001) and time (χ^2^=33.96, df=1, p<0.001), but no interaction was included. Posthoc pairwise analysis indicated significant differences between tuna and termite baits, regardless of the presence of visual cues **(Table S1)**. No significant difference was found in the time when ants first entered the experimental tubes (χ^2^=1.3062, df=3, p=0.728) **(Fig. 3b)**.

### 3.4 Olfactory and visual (caterpillar shape) cues

Aligning with the Olfaction and natural shape experiment, plasticine cubes with tuna paste (4.19±1.48) and plasticine caterpillars with tuna paste (3.58±1.26) attracted more ants than plasticine caterpillars with termite paste (1.75±0.64) and plasticine cubes with termite paste (1.63±0.61) **(Fig. 2c)**. The best model included bait type (χ^2^=39.34, df=3, p<0.001) and time (χ^2^=23.75, df=1, p<0.001), but not their interaction. Posthoc pairwise comparisons showed significant difference between tuna and termite regardless of the plasticine shape **(Table S1)**. Time was also a significant factor in the model showing the overall increase in recruitment of ants, especially to the tuna baits. No significant difference was found in the time when an ant first entered the experimental tubes (χ^2^=0.83, df=3, p=0.842) **(Fig. 3c)**.

### 3.5 Absence of visual cues

The darkroom experiment showed a consistent pattern of the preference for tuna baits, with the largest number of ants visiting tuna cubes (5.03±3.23) followed by tuna caterpillars (3.50±2.26), termite cubes (1.90±1.25), and termite caterpillars (1.87±1.23) **(Fig. 2d)**. Both treatment (χ^2^=31.90, df=3, p<0.001) and time (χ^2^=38.52, df=1, p<0.001) were included in the final model, but not their interaction.

## 4. Discussion

Currently, only two species are in the weaver ant genus *Oecophylla*, one in Oriental and Indo-Australian regions (*Oecophylla smaragdina*) and the other in the Afrotropics (*O. longinoda*), have been described (Boudinot & al. 2022). As the taxonomy of *Oecophylla* has not been well investigated (Garcia & al. 2013), we constructed phylogenetic trees of ants used for our experiments to confirm that all of the ant nests were *Oecophylla smaragdina* from the same clade. In addition, we checked that the relative eye size did not differ among the nests from the two distinct habitats, indicating that the differences in the two habitats (rainforest and rubber plantations) have no evidence of diverging genetically or morphologically in the size of the eye. Taken together, our results are likely to be applicable to *O. smaragdina* of Oriental and Indo-Australian regions found across different habitats.

We used an experimental approach to test for (1) the efficacy of plasticine caterpillar to assess predation and (2) the importance of static visual and olfactory cues for foraging *O. smaragdina*. To our surprise, the ants did not show a preference for plasticine caterpillars compared with plasticine cubes. Both plasticine caterpillars and cubes were hardly visited or attacked by *O. smaragdina* ants. Ants showed a strong preference for tuna compared with termites regardless of the presence of the visual cues, suggesting that static visual cues are not a decisive factor for prey selection. It was suggested in several studies that arboreal ants have evolved to rely more on visual cues and less on olfaction compared with ground-dwelling ants (Jaffe & Perez 1989, Short 2020), however, their conclusions are drawn predominately on neural anatomy while behavioral tests were lacking. Our results suggest that *O. smaragdina*, one of the dominant arboreal ants in this region, relies on olfactory cues for prey selection. Studies with similar behavioral trials on other ant species in different habitats are needed to further elaborate on the relative importance of sensory cues across different habitats and behavioral contexts.

We found a clear and consistent preference for tuna over termite baits, although both baits were taken by *O. smaragdina* workers. *Oecophylla smaragdina* was used as an efficient biological control agent against termites (Musyafa & al. 2019), and during the present study, some individuals were observed to carry the untreated termites out from the tube, demonstrating that the ants do forage on termites. On the other hand, tuna has been commonly used as a bait for *O. smaragdina* and has been shown to be a preferred bait because of its high protein content, and this preference is consistent in the present study (Lach & Hoffmann 2011, Pimid & al. 2019). While tuna as an artificial bait might be a supernormal olfactory stimulus in attracting ants over natural termites, trials with or without the presence of olfaction cues in baits leads to the same conclusion that the ants did not prefer caterpillar shape or termite shape over cube. Additionally, the preference for tuna over termite baits may be attributable to the lack of motion cues in our termite baits. One study (Paluh & al. 2014) showed that sentinel prey with motion increased predation, and this may be the case with *O. smaragdina* as well.

The use of caterpillar-shaped plasticine sentinel models is a well-established experimental approach widely used to study predation pressures at various spatial (elevation, latitude) and temporal (seasons, time of the day) scales (Molleman & al. 2016, Pan & al. 2021, Richards & Coley 2007, Roslin & al. 2017). Although plasticine lacks olfactory cues of natural prey, other studies have demonstrated that the attack rates of plasticine caterpillars were comparable to that of real caterpillars, making this method a useful way of estimating predation rate (Howe & al. 2009, Sam & al. 2015, Richards & Coley 2007). A recent study conducted by Nimalrathna et al. (2023), however, challenged this notion, as they found the attack rate on plasticine caterpillars was substantially lower than that on real caterpillars. Despite the fact that caterpillars are the prey of *O. smaragdina* in their natural habitat, they may not detect caterpillars (and other prey) using static visual cues but rather, the movement of the caterpillar, odors emitted directly by the caterpillar, or indirectly, by the volatile compounds released from the damaged plants (Vet & Dicke 1992, Paluh & al. 2014). While studies based on the use of sentinel caterpillars suggested an increase in predation with canopy habitat compared with understory (Loiselle & Farji-Brener 2002), the present study suggests that the sentinel plasticine prey was not attacked by dominant arboreal predators such as *O. smaragdina* that was predicted to be more visually oriented. Thus, the results of predation studies using sentinel caterpillars miss out on a non-negligible portion of predators that need more than immobile visual cues.

## 6. Acknowledgements

We thank Kyle Tomlinson and Damian Elias for valuable discussion, Zhizhou Jia for termite collection, Ting Tang, Haoran Su, Xianhui Shen, Jingxin Liu and Xue Xia for technical support. This study was supported by the Advanced Field Course in Ecology and Conservation, Xishuangbanna Tropical Botanical Garden, Chinese Academy of Sciences. AN was supported by the National Natural Science Foundation of China International (Regional) Cooperation and Exchange Project (32161160324), Strategic Priority Research Program of Chinese Academy of Sciences (XDB31000000), and the High-End Foreign Experts Program of the High-Level Talent Recruitment Plan of Yunnan Province.

## 8. Author Contributions

AN, LY, SPK, and YL conceived the ideas and designed the methodology; LY, SPK,YL and TM collected the data; LY, SPK and TM analysed the data; LY and AN led the writing of the manuscript. All authors contributed critically to the drafts and gave final approval for publication.

## 11 Supporting information

### 11.1 DNA sample preparation

DNA extraction was conducted by EasyPure® Genomic DNA Kit (TransGen Biotech Co., Ltd., China). CO1 genes were amplified using the primer set LCO1490 (GGTCAACAAATCATAAAGATATTGG) and HCO2198 (TAAACTTCAGGGTGACCAAAAAATCA). For the PCR master mix, 2 µl dNTPs (final 0.2 mM), 2.5 µl 10 ×TransTaq HiFi buffer I, 0.5 µl forward and reverse primers (final 0.2 mM), 0.3 µl TransTaq HiFi DNA polymerase (TransGen Biotech Co., Ltd., China), and 0.2 µl of DNA template were mixed in nuclease-free water, totaling 25 μl per reaction. The PCR reaction was run in a Mastercycler® nexus gradient (Eppendorf, Canada). We employed the touchdown protocol: 2 min of Taq DNA polymerase activation at 95°C; for the first 5 cycles, denaturation at 95°C for 15 seconds; annealing at decreasing temperature from 54°C to 50°C (1°C decrease per cycle) for 30 seconds; extension at 72°C for 1 min. The following 35 cycles involved; at 95°C for 15 seconds; at 49°C for 30 seconds; at 72°C for 1 min. The final extension was at 72°C for 7 min. After confirming the PCR amplification on a 1.5% agarose gel, Sanger sequencing was outsourced from Sangon Biotech Co., Ltd. (https://www.sangon.com/).

### 11.2 DNA Extraction and Phylogenetic Analysis

A total of five worker ants, each from two nests collected from the rainforest and all three nests from the rubber plantation was randomly selected for genetic analysis to confirm our species identification. Ant legs were used for DNA extractions and the mitochondria cytochrome oxidase subunit 1 (COI) gene used for DNA barcoding. Additionally, we incorporated COI sequences of four *Oecophylla smaragdina* workers collected locally (*BR5GB12d201910* and *BR4TB5n201910* collected in the rubber plantation of Bubeng, and *F5TB1d2018* and *F4GB6d2018* collected in the rainforest near XTBG), five *Oecophylla smaragdina* collected from several countries (available from GenBank: *JQ681064; MH686448; HM394876; MW056458; KC685044*), and one *Oecophylla longinoda* (*MT152291*) as an outgroup. The details of the DNA extraction, amplification and sequences are provided in the Supplementary Information.

### 11.3 Relative eye size measurement

To measure relative eye size, we randomly selected six major and six minor workers from each colony. We collected three workers on the nest and another three foraging away from the nest, to account for the potential differences in the colony division of labor. We included both major and minor workers to check whether the differences in relative eye size were driven by the worker castes. All 72 collected individuals were euthanized in a freezer and their heads were removed for imaging under a stereomicroscope (Leica DMC 4500). Eye and head area was then measured from images using ImageJ. Relative eye size was calculated by eye area divided by the head area of individual ants.

**Supplementary Figure S1.**
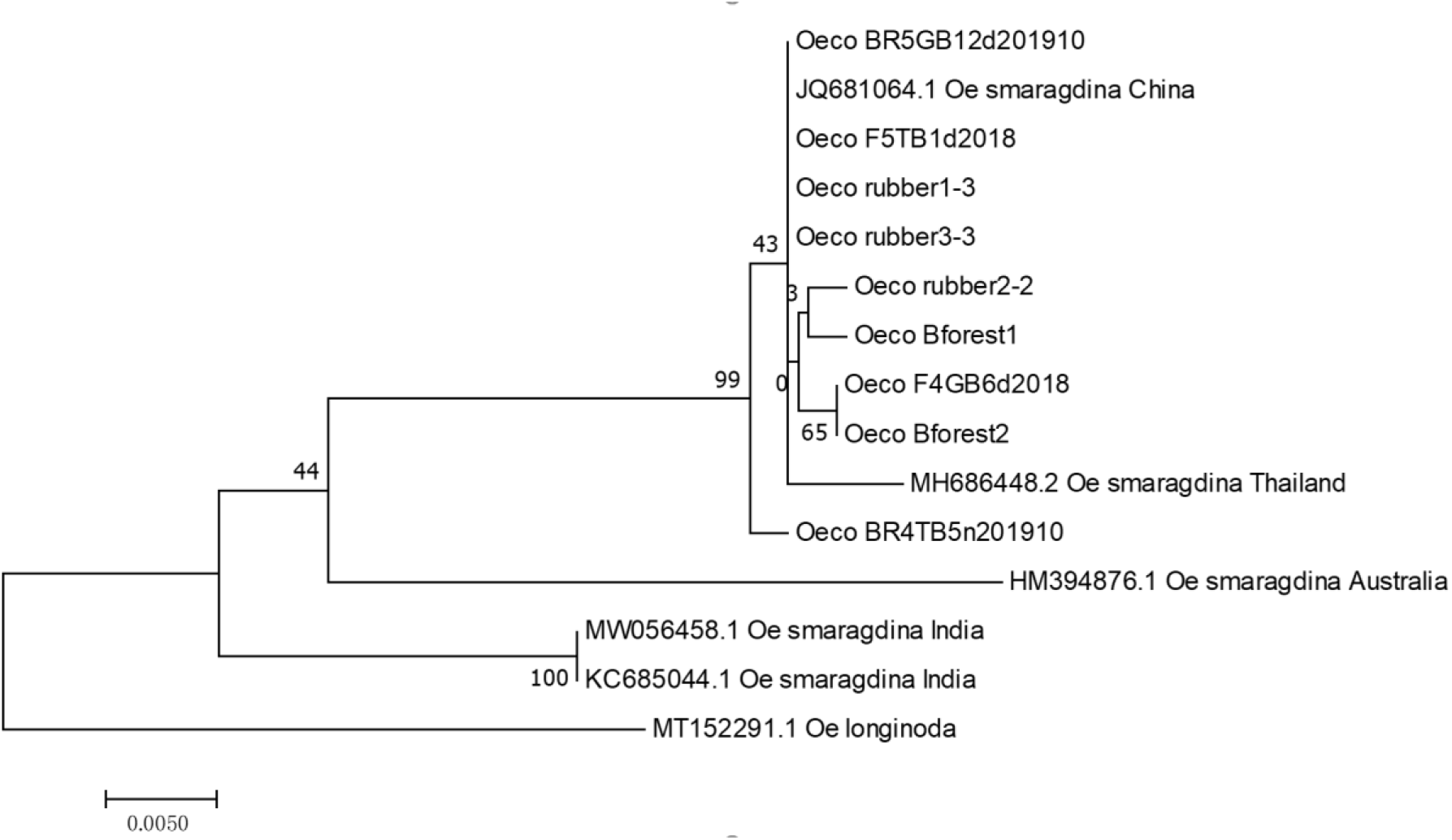
Molecular phylogenetic tree constructed by the maximum likelihood method based on the Jukes-Cantor model (Jukes & Cantor no date). “Oeco rubber” corresponds to the three worker ants each from the three nests we collected from the rubber plantation, and “Oeco Bforest” corresponds to the two worker ants collected from the rainforest canopies. The percentage of trees in which the associated taxa clustered together is shown next to the branches. Initial tree(s) for the heuristic search were obtained automatically by applying the neighbor-join and BioNJ algorithms to the matrix of pairwise distances estimated using the maximum composite likelihood approach, and then selecting the topology with superior log likelihood values. The tree is drawn to scale, with branch lengths measured in the number of substitutions per site. The analysis involved 15 nucleotide sequences. Codon positions included were 1st+2nd+3rd+noncoding. All positions containing gaps and missing data were eliminated.

There was a total of 581 positions in the final dataset. Evolutionary analyses were conducted in MEGA7 (Kumar & al. 2016).

**Supplementary Figure S2.**
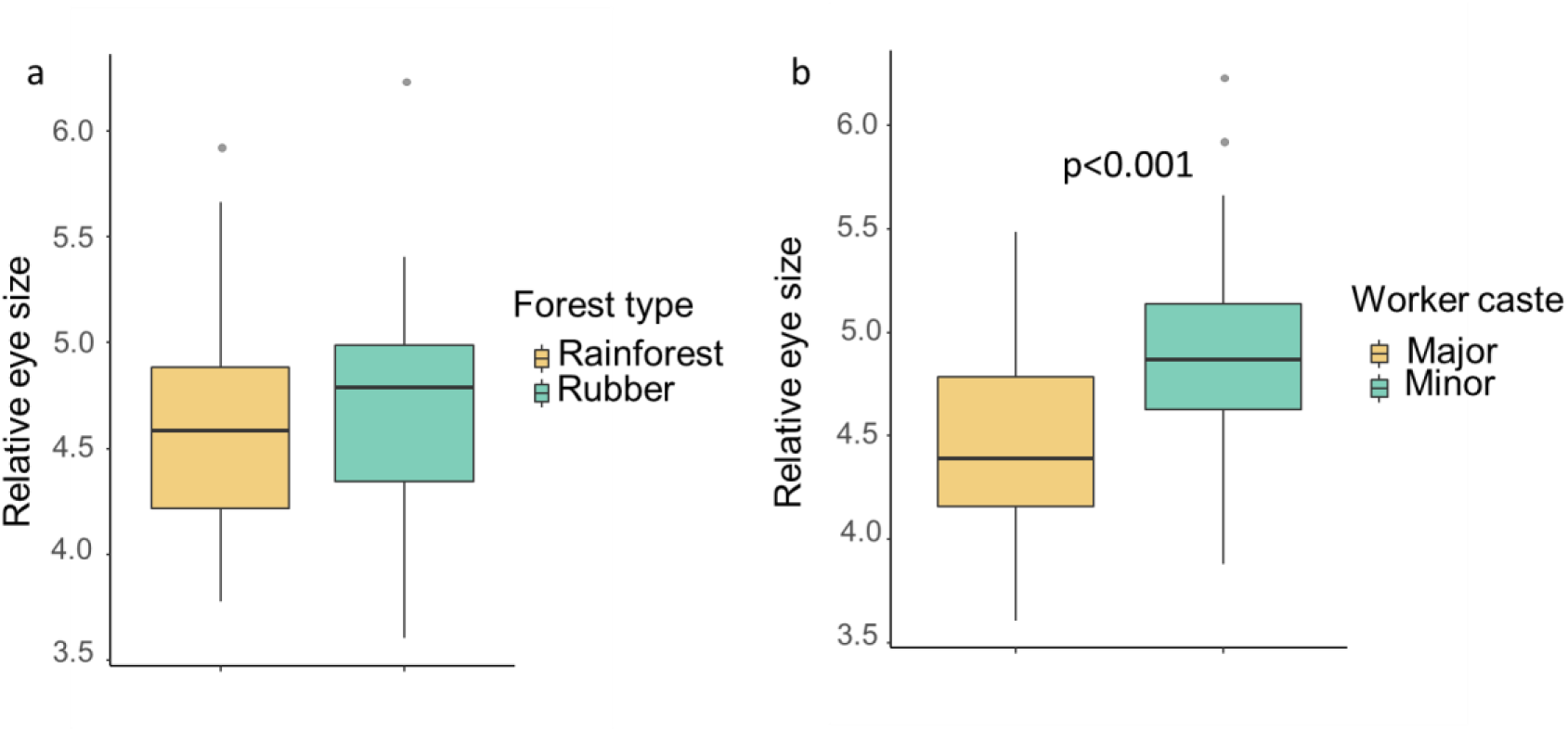
A: relative eye size of ants collected from rainforest and rubber plantations. B: relative eye size of major worker and minor worker. No statistically significant differences were found in relative eye size between forest types that ant nests. Minor workers were found to have significantly larger eyes than major workers (p < 0.001).

**Supplementary Table S1.**
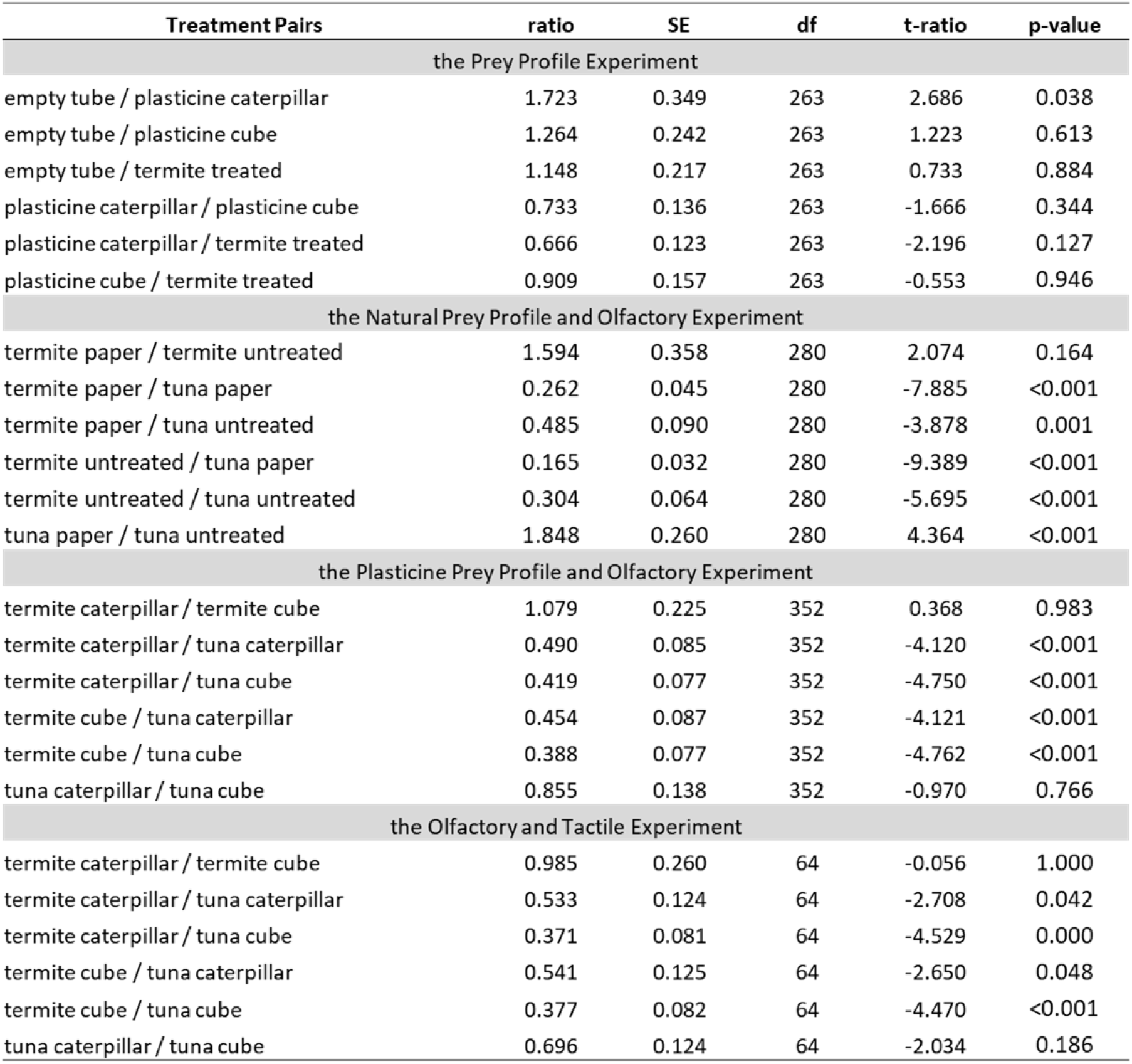
the Output of Tukey’s Honest Significant Difference (HSD) test that compared the treatment pairs.

## Reference

Andersen, A.N. 1992: Regulation of “Momentary” Diversity by Dominant Species in Exceptionally Rich Ant Communities of the Australian Seasonal Tropics. – The American naturalist 140: 401–420.

Anton, S. & Gnatzy, W. 1998: Journal of insect behavior 11: 671–690.

Arnold, T.W. 2010: Uninformative Parameters and Model Selection Using Akaike’s Information Criterion. – The Journal of wildlife management 74: 1175–1178.

Bates, D., Sarkar, D., Bates, M.D. & Matrix, L. 2007: The lme4 package. – R package version 2: 74.

Bolker, B. 2016: Getting started with the glmmTMB package. – Vienna, Austria: R Foundation for Statistical Computing. Software

Boudinot, B.E., Borowiec, M.L. & Prebus, M.M. 2022: Phylogeny, evolution, and classification of the ant genus Lasius, the tribe Lasiini and the subfamily Formicinae (Hymenoptera: Formicidae). – Systematic entomology 47: 113–151.

Brooks, M.E., Kristensen, K., Benthem, K.J. van Magnusson, A., Berg, C.W., Nielsen, A., Skaug, H.J., Mächler, M. & Bolker, B.M. 2017: Modeling zero-inflated count data with glmmTMB. – bioRxiv: 132753.

Brooks, M.E., Kristensen, K., Van Benthem, K.J., Magnusson, A., Berg, C.W., Nielsen, A., Skaug, H.J., Machler, M. & Bolker, B.M. 2017: glmmTMB balances speed and flexibility among packages for zero-inflated generalized linear mixed modeling. – The R journal 9: 378–400.

Brunner, G. 2010: Applications of supercritical fluids. – Annual review of chemical and biomolecular engineering 1: 321–342.

Crawley, M.J. 2009: Natural Enemies: The Population Biology of Predators, Parasites and Diseases. – John Wiley & Sons, 592 pp.

Denan, N., Norhisham, A.R., Sanusi, R., Stone, J. & Azhar, B. 2023: Stand-level habitat characteristics and edge habitats drive biological pest control services in the understory of oil palm plantations. – Biological control: theory and applications in pest management 183: 105261.

Finke, D.L. & Denno, R.F. 2005: Predator diversity and the functioning of ecosystems: the role of intraguild predation in dampening trophic cascades. – Ecology letters 8: 1299–1306.

Finke, D.L. & Denno, R.F. 2004: Predator diversity dampens trophic cascades. – Nature 429: 407–410.

Floren, A., Biun, A. & Linsenmair, E.K. 2002: Arboreal ants as key predators in tropical lowland rainforest trees. – Oecologia 131: 137–144.

Forbes, S.J. & Northfield, T.D. 2017: *Oecophylla smaragdina* ants provide pest control in Australian cacao. – Biotropica 49: 328–336.

Fox, J. & Weisberg, S. 2018: An R Companion to Applied Regression. – SAGE Publications, 608 pp.

Friend, T. 1995: Video techniques in animal ecology and behaviour. – Applied Animal Behaviour Science 43: 303–304.

Garcia, F.H., Wiesel, E. & Fischer, G. 2013: The Ants of Kenya (Hymenoptera: Formicidae)—Faunal Overview, First Species Checklist, Bibliography, Accounts for All Genera, and Discussion on Taxonomy and Zoogeography. – Journal of East African Natural History 101: 127–222.

Hardin, J.W., Hardin, J.W., Hilbe, J.M. & Hilbe, J. 2007: Generalized Linear Models and Extensions, Second Edition. – Stata Press, 387 pp.

Howe, A., Lövei, G.L. & Nachman, G. 2009: Dummy caterpillars as a simple method to assess predation rates on invertebrates in a tropical agroecosystem. – Entomologia experimentalis et applicata 131: 325–329.

Jackson, R.R. & Pollard, S.D. 1996: Predatory behavior of jumping spiders. – Annual review of entomology 41: 287–308.

Jaffe, K. & Perez, E. 1989: Comparative study of brain morphology in ants. – Brain, behavior and evolution 33: 25–33.

Jeanne, R.L. 1979: A Latitudinal Gradient in Rates of Ant Predation. – Ecology 60: 1211.

Jukes, T.H. and Cantor, C.R. 1969 Evolution of Protein Molecules. – Munro, H.N., Ed., Mammalian Protein Metabolism, Academic Press, New York, 21–132.

Kielty, J.P., Allen-Williams, L.J., Underwood, N. & Eastwood, E.A. 1996: Behavioral responses of three species of ground beetle (Coleoptera: Carabidae) to olfactory cues associated with prey and habitat. – Journal of insect behavior 9: 237–250.

Klärner, D. & Barth, F.G. 1982: Vibratory signals and prey capture in orb-weaving spiders (Zygiella x-notata, Nephila clavipes; Araneidae). – Journal of comparative physiology. A, Neuroethology, sensory, neural, and behavioral physiology 148: 445–455.

Kumar, S., Stecher, G. & Tamura, K. 2016: MEGA7: Molecular Evolutionary Genetics Analysis Version 7.0 for Bigger Datasets. – Molecular biology and evolution 33: 1870–1874.

Lach, L. & Hoffmann, B.D. 2011: Are invasive ants better plant-defense mutualists? A comparison of foliage patrolling and herbivory in sites with invasive yellow crazy ants and native weaver ants. – Oikos 120: 9–16.

Leles, B., Xiao, X., Pasion, B.O., Nakamura, A. & Tomlinson, K.W. 2017: Does plant diversity increase top-down control of herbivorous insects in tropical forest? – Oikos 126: 1142–1149.

Lima, S.L. 1998: Nonlethal Effects in the Ecology of Predator-Prey Interactions. – Bioscience 48: 25–34.

Liu, X., Wang, Z., Huang, C., Li, M., Bibi, F., Zhou, S. & Nakamura, A. 2020: Ant assemblage composition explains high predation pressure on artificial caterpillars during early night. – Ecological entomology 45: 547–554.

Loiselle, B.A. & Farji-Brener, A.G. 2002: What’s up? An Experimental Comparison of Predation Levels between Canopy and Understory in a Tropical Wet Forest. – Biotropica 34: 327–330.

Lokkers, C. 1990: Colony dynamics of the green tree ant (Oecophylla smaragdina Fab.) in a seasonal tropical climate. – Doctoral dissertation, James Cook University.

Maheshwari, P., Ooi, E.T. & Nikolov, Z.L. 1995: Off-flavor removal from soy-protein isolate by using liquid and supercritical carbon dioxide. – Journal of the American Oil Chemists’ Society 72: 1107–1115.

Mishra, M. & Bhadani, S. 2017: Daily activity and visual discrimination reflects the eye organization of weaver ant *Oecophylla smaragdina* (Insecta: Hymenoptera: Formicidae). – bioRxiv, p.193243.

Molleman, F., Remmel, T. & Sam, K. 2016: Phenology of Predation on Insects in a Tropical Forest: Temporal Variation in Attack Rate on Dummy Caterpillars. – Biotropica 48: 229–236.

Musyafa Hasan Bahri, S. & Supriyo, H. 2019: Potential of Weaver Ant (Oecophylla smaragdina Fabricius, 1775) as Biocontrol Agent for Pest of Teak Stand in Wanagama Forest, Gunungkidul, Yogyakarta, Indonesia. – KnE Life Sciences: 239–244.

Narendra, A., Ali, T.M. & Series, S.C. 2012: Ant Species Composition and Diversity in the Sharavathi River Basin, Central Western Ghats. – Sahyadri Conserv. Ser. 3.

Nimalrathna, T.S., Solina, I.D., Mon, A.M., Pomoim, N., Bhadra, S., Zvereva, E.L., Sam, K. & Nakamura, A. 2023: Estimating predation pressure in ecological studies: controlling bias imposed by using sentinel plasticine prey. – Entomologia experimentalis et applicata 171: 56–67.

Oudenhove, L. van Billoir, E., Boulay, R., Bernstein, C. & Cerdá, X. 2011: Temperature limits trail following behaviour through pheromone decay in ants. – Die Naturwissenschaften 98: 1009–1017.

Paluh, D.J., Hantak, M.M. & Saporito, R.A. 2014: A Test of Aposematism in the Dendrobatid Poison Frog Oophaga pumilio: The Importance of Movement in Clay Model Experiments. – Journal of herpetology 48: 249–254.

Pan, X., Mizuno, T., Ito, K., Ohsugi, T., Nishimichi, S., Nomiya, R., Ohno, M., Yamawo, A. & Nakamura, A. 2021: Assessing temporal dynamics of predation and effectiveness of caterpillar visual defense using sawfly larval color and resting posture as a model. – Insect science 28: 1800–1815.

Peckarsky, B.L., Abrams, P.A., Bolnick, D.I., Dill, L.M., Grabowski, J.H., Luttbeg, B., Orrock, J.L., Peacor, S.D., Preisser, E.L., Schmitz, O.J. & Trussell, G.C. 2008: REVISITING THE CLASSICS: CONSIDERING NONCONSUMPTIVE EFFECTS IN TEXTBOOK EXAMPLES OF PREDATOR–PREY INTERACTIONS. – Ecology 89: 2416–2425.

Preisser, E.L., Orrock, J.L. & Schmitz, O.J. 2007: Predator hunting mode and habitat domain alter nonconsumptive effects in predator-prey interactions. – Ecology 88: 2744–2751.

Richards, L.A. & Coley, P.D. 2007: Seasonal and habitat differences affect the impact of food and predation on herbivores: a comparison between gaps and understory of a tropical forest. – Oikos 116: 31–40.

Roslin, T., Hardwick, B., Novotny, V., Petry, W.K., Andrew, N.R., Asmus, A., Barrio, I.C., Basset, Y., Boesing, A.L., Bonebrake, T.C., Cameron, E.K., Dáttilo, W., Donoso, D.A., Drozd, P., Gray, C.L., Hik, D.S., Hill, S.J., Hopkins, T., Huang, S., Koane, B., Laird-Hopkins, B., Laukkanen, L., Lewis, O.T., Milne, S., Mwesige, I., Nakamura, A., Nell, C.S., Nichols, E., Prokurat, A., Sam, K., Schmidt, N.M., Slade, A., Slade, V., Suchanková, A., Teder, T., Nouhuys, S. van Vandvik, V., Weissflog, A., Zhukovich, V. & Slade, E.M. 2017: Higher predation risk for insect prey at low latitudes and elevations. – Science 356: 742–744.

Rößler, D.C., Pröhl, H. & Lötters, S. 2018: The future of clay model studies. – BMC Zoology 3: 1–5.

Sam, K., Remmel, T. & Molleman, F. 2015: Material affects attack rates on dummy caterpillars in tropical forest where arthropod predators dominate: an experiment using clay and dough dummies with green colourants on various plant species. – Entomologia experimentalis et applicata 157: 317–324.

Schmitz, O.J., Hambäck, P.A. & Beckerman, A.P. 2000: Trophic Cascades in Terrestrial Systems: A Review of the Effects of Carnivore Removals on Plants. – The American naturalist 155: 141–153.

Schmitz, O.J., Krivan, V. & Ovadia, O. 2004: Trophic cascades: the primacy of trait-mediated indirect interactions. – Ecology letters 7: 153–163.

Pimid, M., Ahmad, A.H., Krishnan, K.T. and Scian, J., 2019. Food preferences and foraging activity of asian weaver ants, Oecophylla smaragdina (Fabricius)(Hymenoptera: Formicidae). – Tropical Life Sciences Research, 30(2), pp.167–179.

Searle, S.R., Speed, F.M. & Milliken, G.A. 1980: Population Marginal Means in the Linear Model: An Alternative to Least Squares Means. – The American statistician 34: 216–221.

Short, H.E. 2020: Olfactory tissue investment in tropical rainforest ants scales with body size. – : 35.

Tiede, Y., Schlautmann, J., Donoso, D.A., Wallis, C.I.B., Bendix, J., Brandl, R. & Farwig, N. 2017: Ants as indicators of environmental change and ecosystem processes. – Ecological indicators 83: 527–537.

Tsuji, K., Hasyim, A., Harlion & Nakamura, K. 2004: Asian weaver ants, Oecophylla smaragdina, and their repelling of pollinators. – Ecological research 19: 669–673.

Tvardikova, K. & Novotny, V. 2012: Predation on exposed and leaf-rolling artificial caterpillars in tropical forests of Papua New Guinea. – Journal of tropical ecology 28: 331–341.

Vet, L.E.M. & Dicke, M. 1992: Ecology of Infochemical Use by Natural Enemies in a Tritrophic Context. – Annual review of entomology 37: 141–172.

Ward, Hart, Webster & Atton 2007: Turbidity and foraging rate in threespine sticklebacks: the importance of visual and chemical prey cues. – Behaviour 144: 1347–1360.

Wen, X.-L., Wen, P., Dahlsjö, C.A.L., Sillam-Dussès, D. & Šobotník, J. 2017: Breaking the cipher: ant eavesdropping on the variational trail pheromone of its termite prey. – Proceedings of the Royal Society B: Biological Sciences 284: 20170121.

Zvereva, E.L., Castagneyrol, B., Cornelissen, T., Forsman, A., Hernández-Agüero, J.A., Klemola, T., Paolucci, L., Polo, V., Salinas, N., Theron, K.J., Xu, G., Zverev, V. & Kozlov, M.V. 2019: Opposite latitudinal patterns for bird and arthropod predation revealed in experiments with differently colored artificial prey. – Ecology and evolution 9: 14273–14285.

Zvereva, E.L. & Kozlov, M.V. 2023: Predation risk estimated on live and artificial insect prey follows different patterns. – Ecology 104: e3943.

