## supporting information and supplementary figure S1-S2, supplementary table S1 for "Olfaction foraging in visually oriented tropical arboreal ants *Oecophylla smaragdina*: Implications for insect predation studies using artificial sentinel prey"

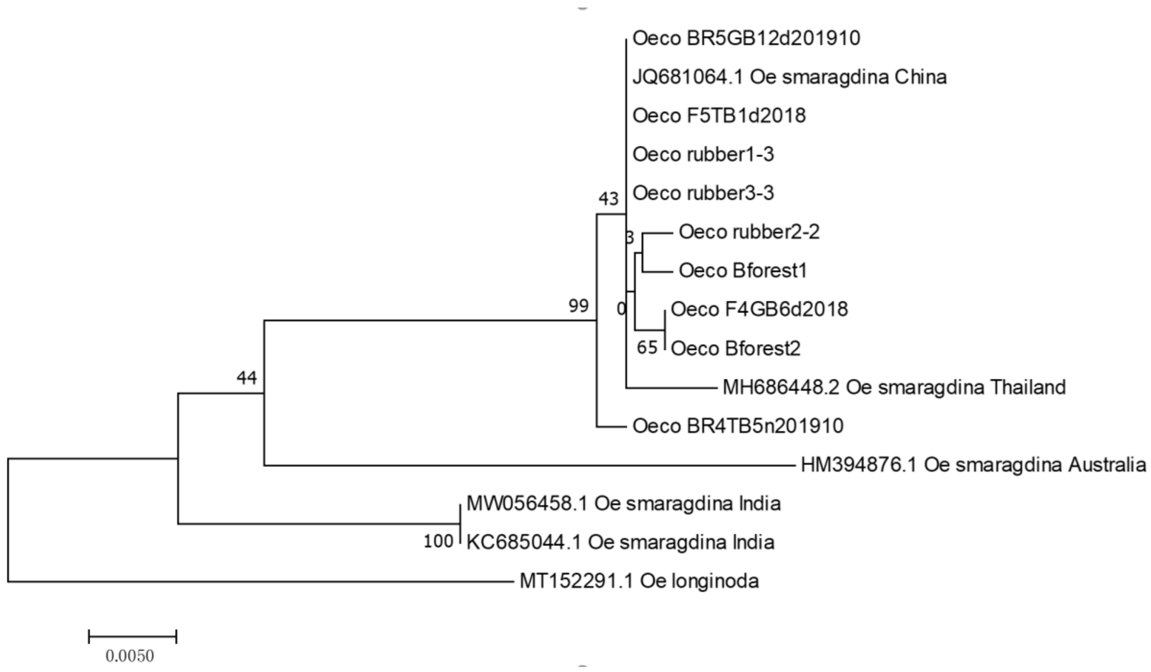


**Supplementary Figure S1** Molecular phylogenetic tree constructed by the maximum likelihood method based on the Jukes-Cantor model (Jukes & Cantor no date). "Oeco rubber" corresponds to the three worker ants each from the three nests we collected from the rubber plantation, and "Oeco Bforest" corresponds to the two worker ants collected from the rainforest canopies. The percentage of trees in which the associated taxa clustered together is shown next to the branches. Initial tree(s) for the heuristic search were obtained automatically by applying the neighbor-join and BioNJ algorithms to the matrix of pairwise distances estimated using the maximum composite likelihood approach, and then selecting the topology with superior log likelihood values. The tree is drawn to scale, with branch lengths measured in the number of substitutions per site. The analysis involved 15 nucleotide sequences. Codon positions included were 1st+2nd+3rd+noncoding. All positions containing gaps and missing data were eliminated.

There was a total of 581 positions in the final dataset. Evolutionary analyses were conducted in MEGA7 (Kumar & al. 2016).

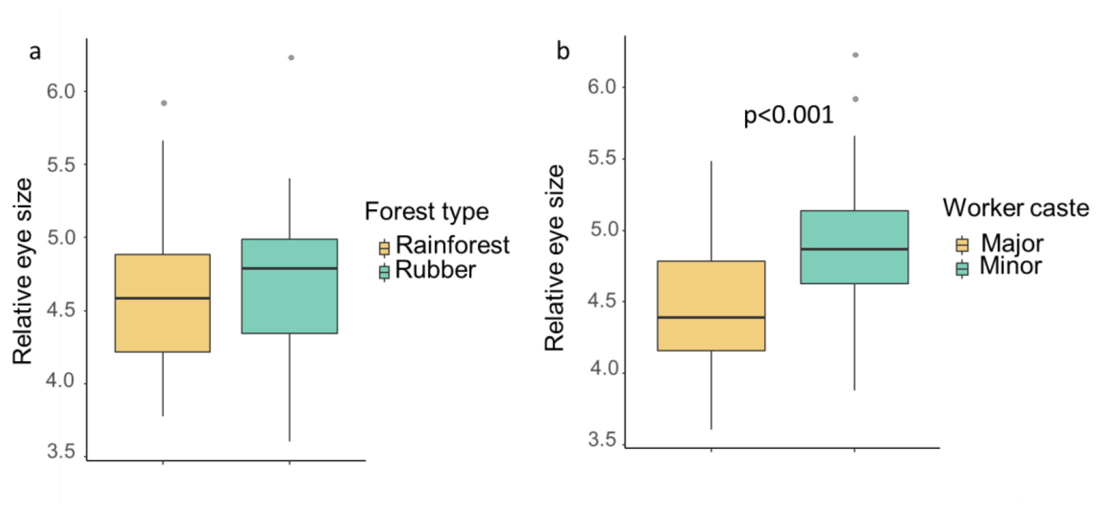


**Supplementary Figure S2**. A: relative eye size of ants collected from rainforest and rubber plantations. B: relative eye size of major worker and minor worker. No statistically significant differences were found in relative eye size between forest types that ant nests. Minor workers were found to have significantly larger eyes than major workers ($p < 0.001$).

**Supplementary Table S1** the Output of Tukey's Honest Significant Difference (HSD) test that compared the treatment pairs.

| Treatment Pairs | ratio | SE | df | t-ratio | p-value |
|---|---|---|---|---|---|
| the Prey Profile Experiment | | | | | |
| empty tube / plasticine caterpillar | 1.723 | 0.349 | 263 | 2.686 | 0.038 |
| empty tube / plasticine cube | 1.264 | 0.242 | 263 | 1.223 | 0.613 |
| empty tube / termite treated | 1.148 | 0.217 | 263 | 0.733 | 0.884 |
| plasticine caterpillar / plasticine cube | 0.733 | 0.136 | 263 | -1.666 | 0.344 |
| plasticine caterpillar / termite treated | 0.666 | 0.123 | 263 | -2.196 | 0.127 |
| plasticine cube / termite treated | 0.909 | 0.157 | 263 | -0.553 | 0.946 |
| the Natural Prey Profile and Olfactory Experiment | | | | | |
| termite paper / termite untreated | 1.594 | 0.358 | 280 | 2.074 | 0.164 |
| termite paper / tuna paper | 0.262 | 0.045 | 280 | -7.885 | <0.001 |
| termite paper / tuna untreated | 0.485 | 0.090 | 280 | -3.878 | 0.001 |
| termite untreated / tuna paper | 0.165 | 0.032 | 280 | -9.389 | <0.001 |
| termite untreated / tuna untreated | 0.304 | 0.064 | 280 | -5.695 | <0.001 |
| tuna paper / tuna untreated | 1.848 | 0.260 | 280 | 4.364 | <0.001 |
| the Plasticine Prey Profile and Olfactory Experiment | | | | | |
| termite caterpillar / termite cube | 1.079 | 0.225 | 352 | 0.368 | 0.983 |
| termite caterpillar / tuna caterpillar | 0.490 | 0.085 | 352 | -4.120 | <0.001 |
| termite caterpillar / tuna cube | 0.419 | 0.077 | 352 | -4.750 | <0.001 |
| termite cube / tuna caterpillar | 0.454 | 0.087 | 352 | -4.121 | <0.001 |
| termite cube / tuna cube | 0.388 | 0.077 | 352 | -4.762 | <0.001 |
| tuna caterpillar / tuna cube | 0.855 | 0.138 | 352 | -0.970 | 0.766 |
| the Olfactory and Tactile Experiment | | | | | |
| termite caterpillar / termite cube | 0.985 | 0.260 | 64 | -0.056 | 1.000 |
| termite caterpillar / tuna caterpillar | 0.533 | 0.124 | 64 | -2.708 | 0.042 |
| termite caterpillar / tuna cube | 0.371 | 0.081 | 64 | -4.529 | 0.000 |
| termite cube / tuna caterpillar | 0.541 | 0.125 | 64 | -2.650 | 0.048 |
| termite cube / tuna cube | 0.377 | 0.082 | 64 | -4.470 | <0.001 |
| tuna caterpillar / tuna cube | 0.696 | 0.124 | 64 | -2.034 | 0.186 |
